# Auxiliary subunits keep AMPA receptors compact during activation and desensitization

**DOI:** 10.1101/295105

**Authors:** Jelena Baranovic, Andrew J.R. Plested

## Abstract

Signal transduction at vertebrate excitatory synapses involves the activity of ionotropic glutamate receptors, including the AMPA (α-amino-3-hydroxy-5-methyl-4-isoxazole propionate) receptor. Technical advances in cryo-electron microscopy have brought a slew of full-length structures of AMPA receptors, on their own and in combination with auxiliary subunits. These structures illustrate a wide range of conformations, indicating that individual domains might undergo substantial lateral motions during gating, resulting in an open, “relaxed” extracellular layer. Here, we used bifunctional methanethiosulfonate cross-linkers to calibrate the conformations found in functional AMPA receptors both in the presence and absence of the auxiliary subunit Stargazin. Our data indicate that AMPA receptors have considerable conformational freedom and can get trapped in stable, relaxed conformations, especially upon long exposures to glutamate. In contrast, Stargazin limits this conformational flexibility. Thus, under synaptic conditions, where brief glutamate exposures and the presence of Stargazin dominate, AMPA receptors are unlikely to adopt very relaxed conformations during gating.

## Introduction

AMPA-type glutamate receptors are found at excitatory synapses throughout the mammalian brain, where they convert glutamate release into membrane depolarisation. Their fast kinetics (Colquhoun et al., 1992; Geiger et al., 1995; Taschenberger and von Gersdorff, 2000), as well as the physical attributes of synapses (Xu-Friedman and Regehr, 2003), allow them to follow glutamate transients at rates above 100 Hz. However, the structural dynamics underlying their rapid signalling are unclear. AMPA receptors are tetrameric ligand-gated ion channels with unique architectural features and loosely coupled structural domains: unstructured linkers connect the amino- and ligand-binding domains (ATDs and LBDs, respectively) forming the extracellular part of the receptor. The LBDs are in turn connected to the transmembrane region (TM) that harbors the integral ion channel (Figure 1A). The extracellular domains adopt local dimer pairs in resting and active receptors. The LBD dimers ‘break up’ in the desensitized state (Sun et al., 2002; Armstrong et al., 2006; Dürr et al., 2014; Twomey et al., 2017b), but motions of the ATDs are unclear (Herguedas et al., 2016; Cais et al., 2014; Yelshanskaya et al., 2016) (Shaikh et al., 2016).

**Figure 1.**
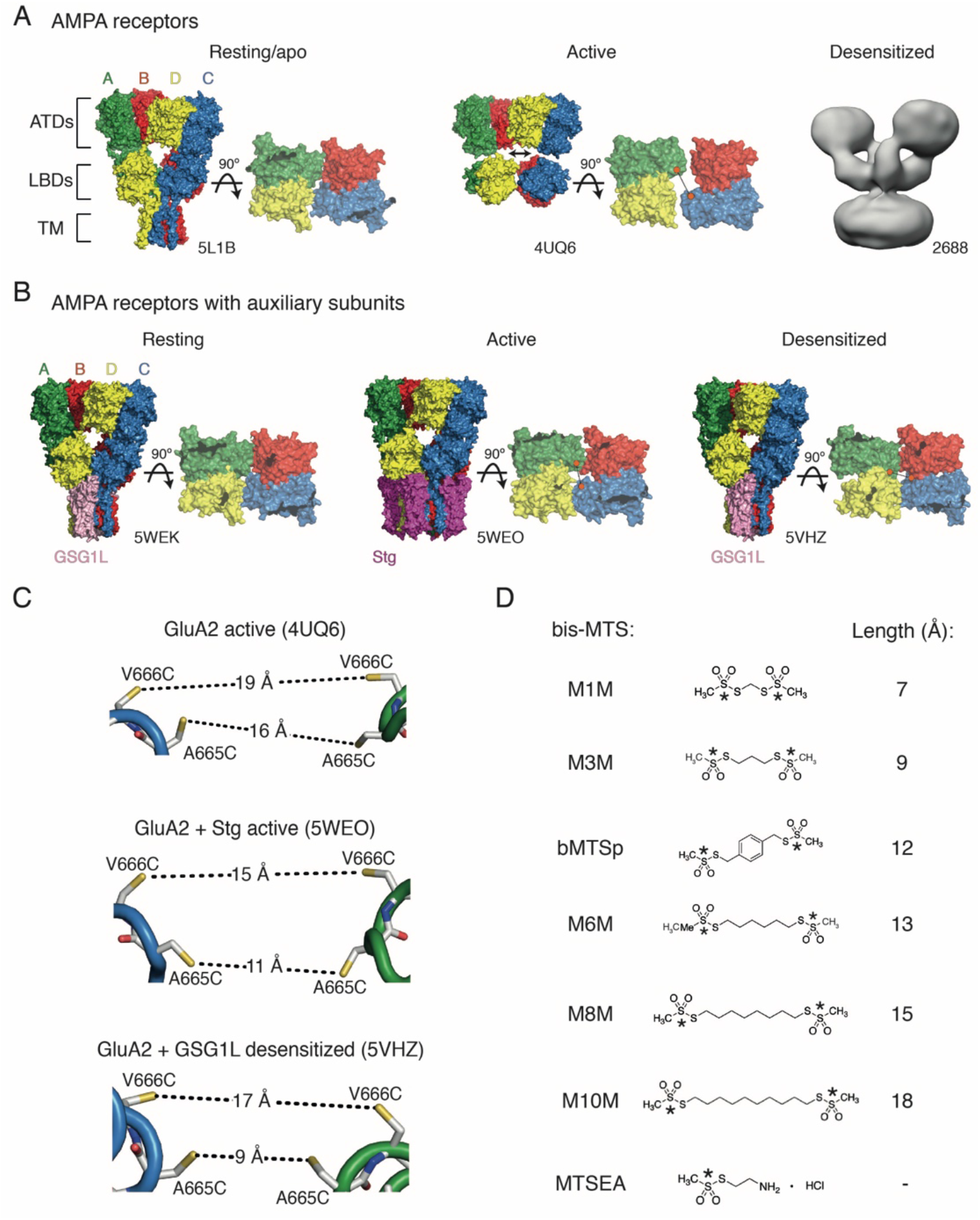
Geometry of AMPA receptors. (A) Structural models of full-length AMPA receptors in resting, active and desensitized states. Accesion codes for PBD or EMDB are indicated. Subunits are colour-coded: A – green, B – red, C – blue and D – yellow. Square brackets delineate AMPA receptor domains: ATDs – amino terminal domains, LBDs – ligand binding domains and TM – transmembrane region. The cytoplasmic domain is not resolved. For the resting and active structures, LBDs are also shown in a top-down view (omitted for the desensitized structure due to its low resolution). Orange spheres connected by black lines indicate 665-666 residues (mutated in this study) in the LBD layer. (B) Same as in (A), but for GluA2 structures complexed with auxiliary subunits: Stg – Stargazin (dark purple) and GSG1L (light purple). (C) Distances between sulfhydryl groups of mutated residues, A665C and V666C, in agonist-bound strucures shown in (A) and (B). (D) Structures of bifunctional (bis-MTS: M1M-M10M) and monofunctional (MTSEA) compounds. Lengths were measured between reactive sulphur atoms (SG, asterisks).

The crowded and narrow synaptic cleft is scarcely wider than the receptors are tall themselves and has narrow edges (Zuber et al., 2005; Tao et al., 2018). This observation implies that conformational dynamics of the receptor domains and their relation to synapse dimensions and molecular composition has implications in both health and disease. For example, if extremely dilated conformations of AMPA receptors can be adopted rapidly, that is on the millisecond timescale of fast excitatory transmission, this could impact receptor anchoring at synapses. Activity-dependent anchoring might be a way to regulate synaptic strength (Constals et al., 2015). On the other hand, slow rearrangements could be relevant for trafficking, and in disease states.

Advances in the structural biology of ionotropic glutamate receptors (iGluRs) have produced a catalogue of static conformational snapshots. Very similar structures have been obtained in conditions corresponding to nominally different functional states (Chen et al., 2014; Dürr et al., 2014; Yelshanskaya et al., 2014). In addition, several structures suggest substantial movements of the extracellular domains when bound by agonist (Nakagawa et al., 2005; Dürr et al., 2014; Meyerson et al., 2014). In these structures, the domains “fall apart”, either breaking local symmetry states, adopting higher order symmetries or switching into a ring-like arrangement (Figure 1A). The timescale of this broad range of potential lateral movements is unknown, because the structural experiments necessarily took place over hours. We therefore set out to investigate the conformational range of agonist bound AMPA receptors with the aim of distinguishing frequently-visited, short-lived conformations from the long-lived ones that likely have less direct relevance to synaptic transmission.

Previously, we demonstrated disulphide bonds and metal bridges trapping receptors in compact LBD arrangements (Salazar et al., 2017; Baranovic et al., 2016; Lau et al., 2013). To measure the separation of domains in this work, we used bifunctional methanethiosulfonate cross-linkers (bis-MTS) of defined lengths (Loo and Clarke, 2001; Guan et al., 2002; Armstrong et al., 2006; Tajima et al., 2016) (Figure 1D). These cross-linkers show specific combination with two free thiol groups, provided by cysteine residues that we engineered. The reactivity of these probes can in principle report distances, giving them the property of nanometre-scale molecular rulers.

Overall, our results suggest that the more dilated conformation of the receptor, the slower it is to access, but once attained, these conformations are stable. However, auxiliary subunits restrict the conformational ensemble, maintaining more compact arrangements. This kinetic classification suggests AMPA receptors at synapses have similar, compact geometries regardless of their instantaneous gating state.

## Results

### Desensitized AMPA receptors can adopt ‘relaxed’ conformations

The rupture of the LBD intra-dimer interface is a structural hallmark of AMPA receptor desensitization, as shown by biophysical studies based on the structures of isolated dimers of ligand binding domains (Sun et al., 2002; Armstrong et al., 2006). Some cryo-electron microscopy (cryo-EM) structures of full-length receptors suggest that desensitization might involve further rearrangements of the ligand-binding domains, including separation of the two dimers and ‘dilation’ of the entire extracellular layer in the membrane plane (Figure 1A).

We attempted to capture this movement between LBDs with bis-MTS cross-linkers ranging from 7 to 18 Å in length (Figure 1D). If the LBD layer opened up in the horizontal plane upon receptor desensitization, this movement should create access for longer bis-MTS cross-linkers in the inter-dimer space (orange dots in Figure 1A). The same principle applies to the ATDs, but ATD layer is functionally silent (Pasternack et al., 2002) and its cross-linking does not produce measurable changes in the receptor activity (Yelshanskaya et al., 2016). Structural models and fluorescence studies (Shaikh et al., 2016) so far indicate that the ATDs follow movements of the LBD layer (Figure 1A-B). We assume here that the membrane-bound TM region strongly restricts vertical displacement of the LBDs, that is, the movements we are probing are those approximately parallel to the membrane.

We identified positions 665 and 666 (in the FG loop) in homomeric GluA2 receptors as best positioned to follow separation of the LBD dimers (Figure 1A-C). The presumptive geometries of the sulfhydryl groups (SG) for cysteine mutants at these sites, in the agonist bound states of GluA2 receptor (with and without auxiliary subunits) are shown in Figure 1C. The structure of the desensitized GluA2 receptor (EMDB: 2688, Figure 1A) is not detailed enough to measure residue distances, but the equivalent residues are 21 Å apart in homologous kainate receptors (PDB: 5KUF; (Meyerson et al., 2016)).

As shown in Figure 2B and D, bis-MTS cross-linkers M1M to M10M all caused strong reduction of the peak current in V666C mutant when applied in the desensitized state. To quantify this effect, the peak current was measured from the control pulses before (I_peak_ pre-trap, 4 pulses) and after the 1-minute application (trap) of the cross-linker (I_peak_ post-trap, 2nd control pulse after the trap, arrows in Figure 2A-C). For each patch, the ratio of I_peak_ post-trap over I_peak_ pre-trap was determined and plotted as shown in Figure 2D-E.

**Figure 2.**
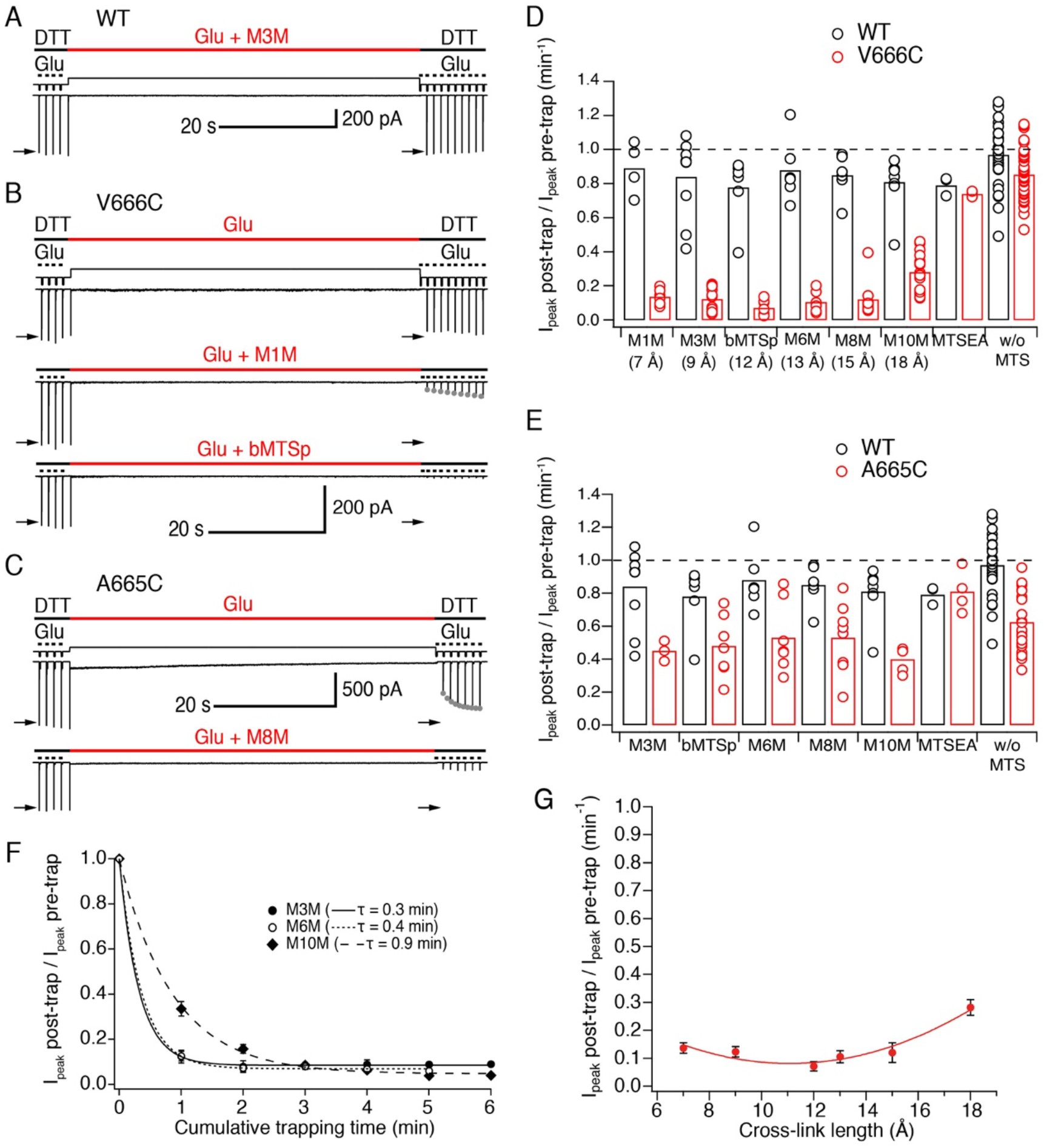
LBDs can separate ≥18 Å in desensitized GluA2. (A) Control recording of wild-type (WT) GluA2 receptors in response to a trapping protocol. Movements of the piezo reflecting the solution exchange are shown in thin, black lines above the current trace. Composition of the solutions is indicated in thick lines above the piezo trace. Downward ticks are 200 ms control jumps from DTT (1 mM) to DTT and glutamate (Glu, 10 mM). Four pre-trap control pulses were followed by a 1-minute long trapping pulse (red line) in Glu (10 mM) and M3M (1 μM). After the trapping pulse, the patch was exposed to 10 post-trap control pulses. The first post-trap pulse gives no response because receptors are desensitized (for details, see Experimental Procedures). (B) Same as in (A), but for V666C mutant. The top two recordings are from the same patch: in the first trace, V666C receptors were exposed only to Glu. Trapping of the same patch in Glu and M1M (1 μM), results in pronounced peak current reduction, which partly recovered with τ_recovery_ = 30 ± 7 s, *n* = 5 (grey dots; post-trap control pulses extended to 30 for fit). The bottom trace is a different patch trapped in bMTSp (1 μM), showing even stronger peak current reduction without any recovery. (C) As in (B) but for the A665C mutant. The two traces are paired recordings of the same patch. Post-trap control pulses show that A665C does cross-link to itself, but most of the current recovers within several seconds after the trap (grey dots; τ_recovery_ = 3.3 ± 0.4 s, n = 17). If the same patch is now trapped in Glu and M8M (1 μM), the peak current reduction is much more pronounced and does not recover. (D) Summary of the trapping effects for WT (black) and V666C (red) for cross-linkers M1M (7 Å) to M10M (18 Å). Trapping effect was calculated as the ratio of the post-trap and pre-trap peak current (arrows in A-C). MTSEA is a monofunctional reagent and “w/o MTS” stands for “without MTS” (traps in Glu only, pooled for all experiments). Dashed line indicates no effect. For peak current reduction in a bis-MTS vs. w/o MTS (pooled), *P* < 10^−7^, for all cross-linkers. (E) Same as in (D), but for A665C (red) in cross-linkers M3M (9 Å) to M10M (18 Å). A665C mutant in the presence and absence of an MTS cross-linker resulted in *P* ≤ 0.02, depending on the cross-linker. For statistics vs. WT and between different bis-MTS, see Table S1. (F) Trapping time for V666C receptors in M3M, M6M and M10M. The 1-min trapping protocol was repeated up to 6 times, resulting in a cumulative exposure of the patch to a bis-MTS of up to 6 min. The data were fit with a monoexponential for each cross-linker (τ indicated in brackets). (G) Trapping profile of desensitized V666C receptors shows the effect of each cross-linker vs. its length, in the first minute of exposure. The data were fit with a parabola (red line): *f*(*x*) = *K*_0_ + *K*_1_^*^(*x* – *K*_2_)^2^ (for details, see Experimental Procedures).

For the V666C mutant, bis-MTS cross-linkers from 7 to 15 Å in length (M1M-M8M; 1 μM), inhibited about 90% of the peak current in the patch after a 1-minute application (see Table S1). The longest (M10M) cross-linker was the slowest one to act (Figure 2F), leading to slightly less inhibition (~70%) in the first minute of exposure. The reduction was less pronounced for A665C mutant for all cross-linkers (~50%, Figure 2E and Table S1).

Is the slow action of bis-MTS cross-linkers (≤ minute, Figure 2F) because we are sampling slowly-attained conformations? We used low concentrations (1 μM) of MTS reagents to ensure bifunctional reagents were not chaining to each other. To determine that modification of the V666C receptors by MTS cross-linkers can proceed on the same time scale as receptor gating we performed additional trapping experiments with 50 μM M3M and M10M (Figure 3). Indeed, at 50 μM, both bis-MTS cross-linkers were roughly 50x faster to modify the receptors (τ_M3M_ = 0.2 s and τ_M10M_ = 1.7 s, Figure 3B and D). The relative modification time was preserved with the longer cross-linker (M10M) still being slower than the shorter one (M3M). These experiments indicate that bifunctional MTS reagents are capturing rapid transitions in receptor structure on the millisecond timescale.

**Figure 3.**
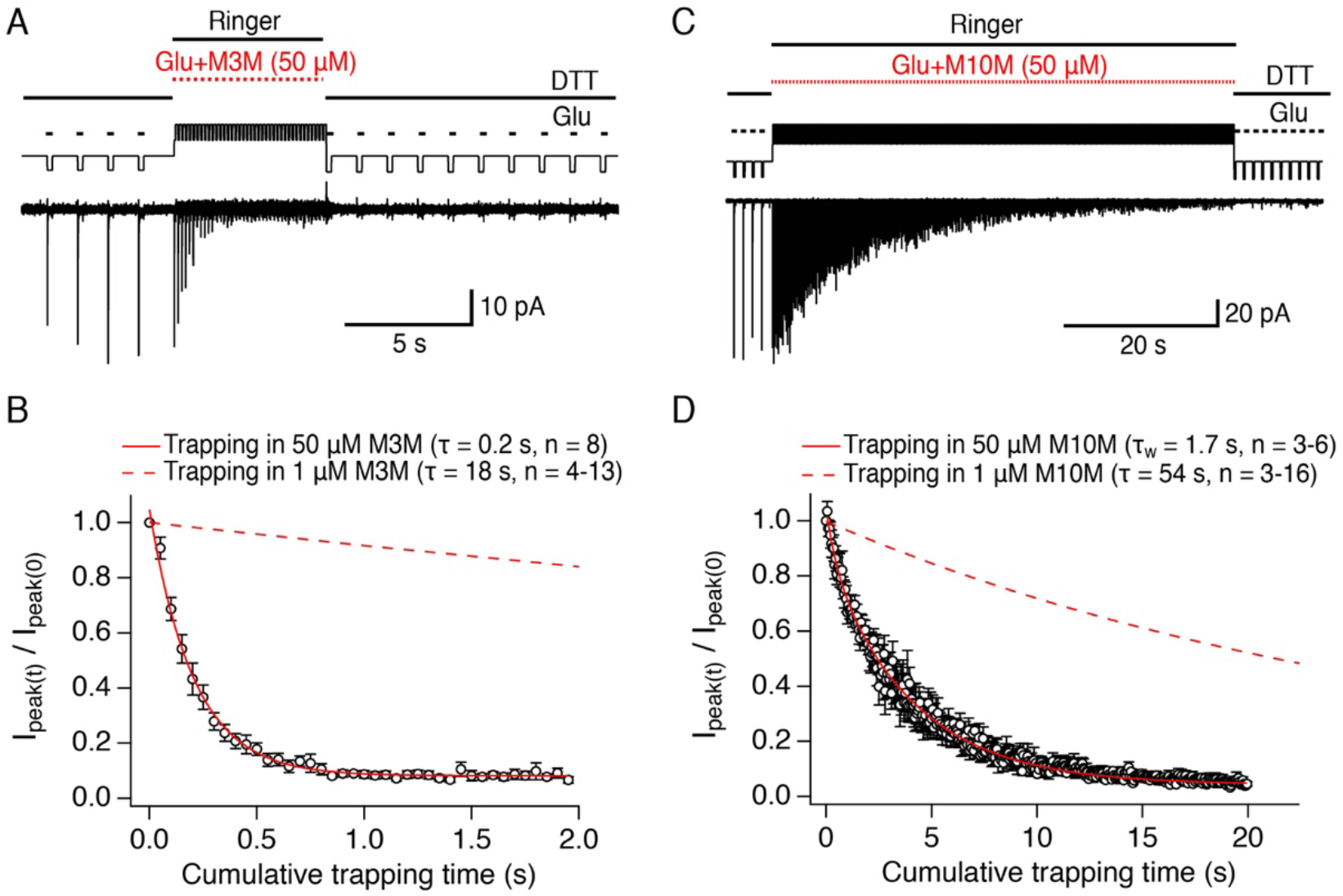
Bis-MTS cross-linkers can trap at millisecond timescale. (A) Trapping protocol for V666C receptors in 50 μM M3M and 10 mM glutamate (Glu), same as in Figure 2A-C, but with a trapping jump consisting of 50 ms-pulses into 50 μM M3M and 10 mM Glu at 6.7 Hz (thick, red lines), interspersed with 100 ms intervals in Ringer solution. (B) Progression of the peak current reduction in response to the trapping pulses (red in (A)) was determined for each patch, normalized to the first pulse, averaged across patches and plotted. The resulting decrease in the peak current was fit with a monoexponential (red line) with τ = 0.2 s. Dashed line is a fit to trapping in 1 μM M3M from Figure 2F. (C) As in (A), but for M10M, with the trapping jump prolonged to 1 minute to complete the peak current reduction. (D) As in (B), but for M10M. Peak current decay was best fit with a double exponential resulting in weighted τ_W_ of 1.7 s.

Inhibition was overall so profound that we sought to establish that it was specific. Two other factors potentially contribute to the current decrease: non-specific run-down of the current and disulphide bonding of the introduced cysteines to each other. Current run-down is particularly difficult to avoid in the long records that we made for these experiments. A665C and V666C sulfhydryl groups were both previously shown to crosslink in the presence of oxidizing agent CuPhen (Salazar et al., 2017; Lau et al., 2013; Yelshanskaya et al., 2016). To account for these confounding factors, we made paired recordings: a patch was first exposed to a 1-minute long trapping pulse containing glutamate only (no cross-linkers), followed by trapping of the same patch in glutamate and a cross-linker (Figure 2B-C). Any run-down in the patch or possible cross-linking of the cysteines to each other was then assessed from trapping in glutamate only. Both mutants underwent some peak current reduction in glutamate in the absence of cross-linker (A665C: 0.62 ± 0.04, *n* = 23, *P* < 10^−7^ vs. WT, V666C: 0.86 ± 0.02, *n* = 44, *P* = 0.003 vs. WT; WT: 0.97 ± 0.03, *n* = 30), consistent with previous results (Lau et al., 2013; Yelshanskaya et al., 2016). Limited cross-linking of A665C cysteines to each other in long exposures to 10 mM glutamate was evident as a recovery of peak responses after the trapping pulse (grey dots in Figure 2C, τ = 3.3 ± 0.4 s, *n* = 17). For the V666C mutant, all 6 bis-MTS cross-linkers (M1M-M10) resulted in more current inhibition than glutamate alone (*P* ≤ 0.03 with and without bis-MTS, depending on the cross-linker; paired randomisation test). Overall, paired recordings gave indistinguishable results to the non-paired recordings. In other words, disulphide crosslinking and rundown were minimal in these conditions. Therefore, we pooled the cross-linking data in glutamate alone across conditions for each mutant (Figure 2D-E).

To confirm that the observed peak current reduction came from crosslinking rather than monofunctional engagement of a cross-linker, we modified mutants with MTSEA (Figure 1D), which can interact only with a single cysteine. As shown in Figure 2D-E, MTSEA failed to inhibit either V666C or A665C above control (I_peak_ post-trap / I_peak_ pre-trap for V666C: 0.89 ± 0.025, *n* = 3-4, *P* = 0.1; for A665C: 0.81 ± 0.06, *n* = 3-4, *P* = 0.8). This result also suggests that the mild oxidizing environment created by MTS compounds at 1 *μ*M has little tendency to promote disulphide bond formation. Thus, bifunctional MTS cross-linking was necessary for the peak current reduction observed in desensitizing AMPA receptors.

The effect of all bis-MTS cross-linkers with respect to their length is summarized in the trapping profile for desensitized V666C receptors (Figure 2G). Rather than showing a preferred cross-linking length (and thus, conformation) for desensitized receptors, the profile is a shallow parabola with strong effects across all tested lengths. Fitting a parabola to these data is justified by the observation that direct disulfide crosslinking (representing the short distance limit) is quite ineffective at this site (Figure 2D and (Lau et al., 2013)). These results demonstrate that desensitized AMPA receptors can ‘open up’ their LBD layer to ~18 Å at position 666 during desensitization and occupy a spectrum of conformations. Structural heterogeneity and ‘dilation’ of the LBD layer to ≥18 Å are both in good agreement with cryo-EM structures (Meyerson et al., 2014; Dürr et al., 2014).

In addition to positions 665 and 666, we also tested nearby positions 662 and 664 for their sensitivity to bis-MTS cross-linkers (Figure S1). The I664C mutant showed similar levels of the peak current reduction to V666C. However, we judged the I664C sulfhydryls were too far apart to make effective use of the available bis-MTS cross-linkers (Figure S1A). Like the A665C mutant, the S662C mutant showed considerable disulphide formation, so we focused on the V666C mutant in further investigations.

Next, we wanted to test how specific the cross-linkers were in targeting the inter-dimer interface of the LBD layer in our conditions. We generated another single cysteine mutant, K493C (Armstrong et al., 2006), positioned within LBD dimers (Figure S2A). Thus, bridging across the two cysteines should keep the LBD dimers intact leading to block of desensitization. When we applied bis-MTS cross-linkers on K493C receptors (1 μM, for ≥1 min), the effects were profoundly different from the mutants at the inter-dimer interface: K493C current underwent potentiation rather than inhibition, with shorter cross-linkers (M1M, M3M and M6M) blocking the receptor desensitization almost completely (Figure S2B-D). We also considered the possibility that bis-MTS cross-linkers might be spuriously cross-linking to wild-type cysteines on the receptor or forming inter-receptor cross-links (Figure S3A). If this were the case, the peak current reduction effect would be expected to scale with the number of the receptors in the membrane (i.e. peak current), but no such correlation was found (Figure S3B). In addition, longer cross-linkers would be expected to be more efficient in forging inter-receptor cross-links, but we found that the longest cross-linker, M10M, was the slowest to react on desensitized AMPA receptors (Figure 2F). The absence of the strong peak current reduction in the WT receptors also speaks against the bis-MTS cross-linkers interacting with native cysteine residues. These results led us to conclude that the bis-MTS cross-linkers are cross-linking cysteines introduced at the LBD inter-dimer interface (V666C).

### Cross-linked desensitized states are highly stable

Upon establishing specific and strong reduction of the peak current in desensitizing V666C receptors by the bis-MTS cross-linkers, we next sought to examine the stability of trapped states. The more stable the trapped state, the longer we would expect that it takes for trapping effects to reverse, and vice versa. The fastest recovery after trapping was observed with M1M (τ = 30 ± 7 s, *n* = 5) and the time constant of the peak current recovery could be measured by directly fitting the peak current of the post-trap control pulses (grey dots in Figure 2B). With longer bis-MTS cross-linkers, the recovery time increased to minutes, making direct measurements of the recovery time from post-trap control pulses impractical. Instead, the experimental design was adjusted to allow measurements of long recovery times as described in Experimental Procedures and Figure 4. After the receptors were trapped with M3M for 1 minute, the peak current in the patch did recover, but very slowly, taking over 10 minutes (600 applications of glutamate) to reach the pre-trapping levels. With longer cross-linkers, the peak current was essentially irreversible over the timescales we could measure (20 minutes; Figure 4D-G). The progressive inability of the receptors to recover from trapping with longer cross-linkers indicates increasingly stable trapped states corresponding to greater separation of the A and C subunits in the desensitized state (Figure 4G-H). Desensitized AMPA receptors with a disulphide bridge formed between the two V666C residues at inter-dimer interface fit in well with this trend, recovering in seconds following 100 s exposure to the oxidizing agent CuPhen (A.P. and Hector Salazar, unpublished data) (Figure 4H).

**Figure 4.**
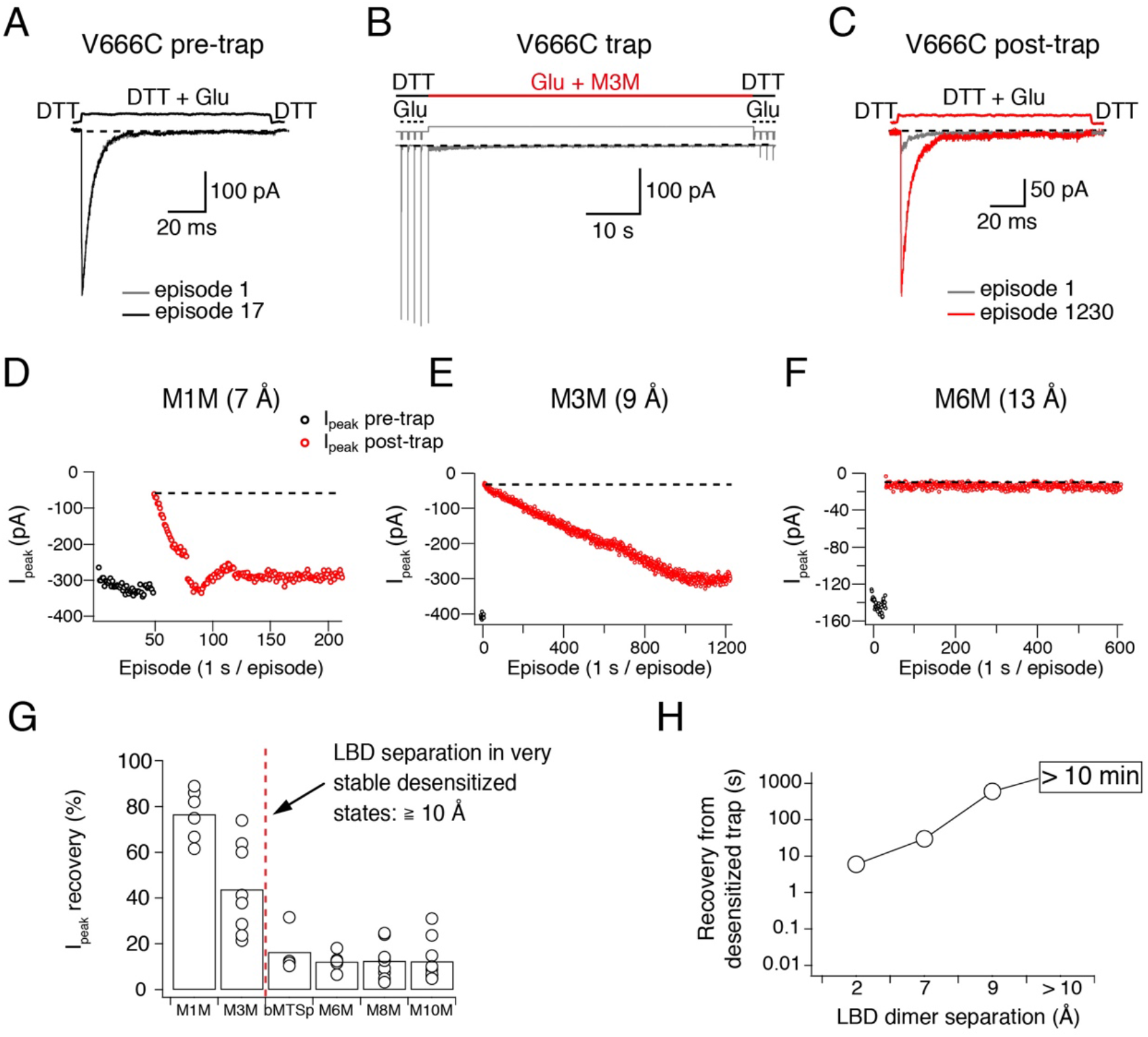
Recovery of trapped desensitized V666C receptors depends on the LBD separation. (A) Protocol to measure recovery from trapping. To obtain a stable baseline response to glutamate, we first repeated brief glutamate applications in reducing conditions (1 mM DTT). In the given example, we gave 17 pulses (100 ms, 1 Hz). (B) In the following step, the patch was exposed to 1 μM cross-linker (here M3M) and 10 mM Glu for 1 minute, with control pulses before and after the trap. (C) After the trapping protocol, the patch was again exposed to fast, reducing glutamate jumps like in (A) in order to follow recovery of the response. In this example, we could record 1230 consecutive episodes (~20 min) and obtain almost complete recovery. Note the difference between the current amplitude in the 1^st^ episode (grey) and the 1230^th^ episode (red). The recovery of the patch current (I_peak_) in typical experiments for different cross-linkers is plotted in panels D - F (panel E is the same patch as in panels A - C). Black dots show the responses before the trap and red dots the peak current after the trap. The gap in red dots in (D) represents a switch between recording protocols. (G) Summary of the peak current recovery for different cross-linkers. The percentage of recovered current is the ratio of the peak currents recorded 3-10 min after the trap to the peak current before trapping. Dashed, red line denotes a limit after which no recovery of the current could be measured within 10 min after the trap. (H) Plot of recovery time from trapped desensitized states vs. the inter-dimer separation at position V666C in the LBD layer. The first data point indicates recovery from V666C disulphide bridges formed in desensitized state (A. P. and Hector Salazar, unpublished data).

To ensure that the lack of recovery was not due to a limited reducing capacity, we tested a higher concentration of the reducing agent DTT (5 mM instead of 1 mM). Stronger reducing conditions did not consistently promote recovery of V666C receptors trapped by a long cross-linker M6M (12 Å; *P* = 0.3, Figure S4).

### Activation limits conformational heterogeneity of the LBD layer

We next investigated if the LBDs of activated V666C receptors are also accessible to a similarly wide range of bis-MTS cross-linkers. According to one cryo-EM structure of an apparently activated AMPA receptor (Meyerson et al., 2014) (Figure 1C), V666C residues should be far enough apart to accommodate cross-linkers up to 19 Å in length.

To maintain the active state, we blocked desensitization with cyclothiazide (CTZ, 100 μM). As shown in Figure 5, block of desensitization reduced inhibition by bis-MTS cross-linkers in the first minute of exposure (see Table S2 for statistics). Current inhibition for every cross-linker in the presence and absence of desensitization is shown in Figure 5B. With M1M, about 15% of V666C receptors recovered from inhibition with a time constant of τ = 7.1 ± 1 s, n = 4, leading to the final inhibition of 0.65 ± 0.02, n = 12 (not shown). For M3M, the recovery was still faster with τ = 1.98 ± 0.2 s, n = 8 (grey dots in Figure 5A), with about 15% the receptors recovering and leading to the final inhibition of 0.53 ± 0.03, n = 9. The fast recovery indicates that with dimer separation of about 9 Å, receptors were trapped in an unstable, stereochemically-strained state. Notably, the SG of V666C are 9 Å apart in a structure solved with the partial agonist NOW, which may represent a pre-open state (Yelshanskaya et al., 2014).

**Figure 5.**
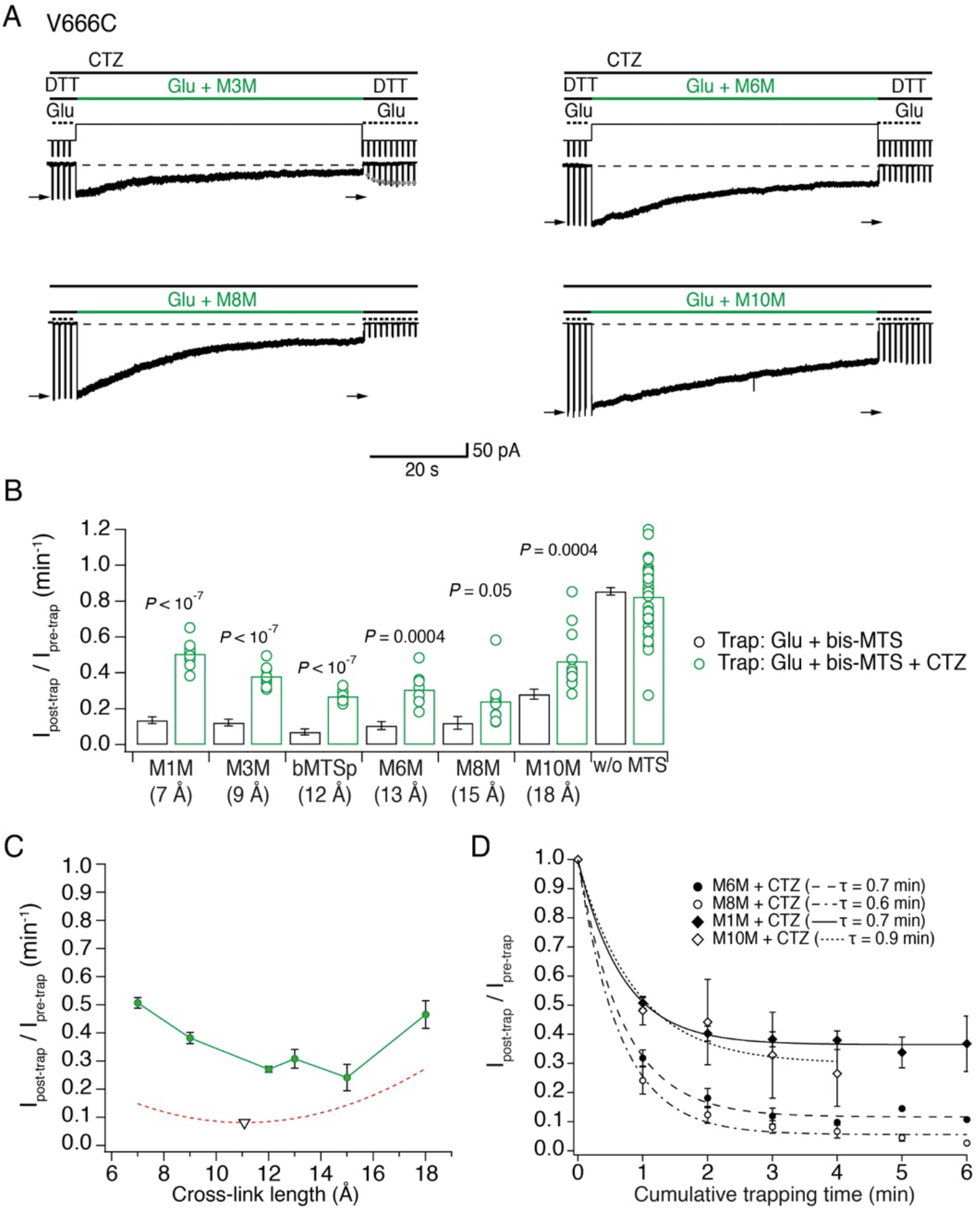
The active LBD layer is dilated. (A) Current traces of trapping protocols on active V666C receptors. Cyclothiazide (CTZ) was included at 100 μM throughout the experiment to block desensitization. The duration of the trapping pulse is indicated in green. Trapping rate (τ) was determined from the monoexponential fits (red, dotted line; *n*: number of patches). For M3M, gray dots indicate recovery from the trap immediately after the trap (τ = 2.0 ± 0.2 s, *n* = 8). Extent of trapping was relative to the initial pulses (arrows) (B) Summary of bis-MTS trapping (1 min) of desensitizing (black) and non-desensitizing V666C receptors (green). Desensitizing data are from Figure 2E. *P* values compare the desensitizing and non-desensitizing condition. For *P* values comparing effects with and without bis-MTS and between the cross-linkers, see Table S2, respectively. Bis-MTS length is indicated in brackets; “w/o MTS”: without MTS. (C) Trapping profile of active V666C receptors after the first minute of exposure (green). Green line connects the points. Fit of the trapping profile of desensitized receptors is shown as a red, dashed line for comparison. Black triangle indicates minimum of the fit at 11 Å. (D) Dependence of the trapping time on the length of the bis-MTS. V666C receptors were exposed up to 6 times to the trapping protocol described in (A). Current reduction was determined after each 1-min application to a bis-MTS. Current decay was described with a monexponential fit (τ in brackets).

With desensitization blocked, AMPA receptors displayed a distinct trapping profile from that of desensitized receptors (Figure 5C). The sharper trapping profile could not be described by a parabola and indicated a favored LBD separation of 15 Å, greater than the shallow optimum of desensitized receptors (11 Å). Because inhibition was less profound than in desensitized receptors for all cross-linkers except M8M, we investigated the possibility that bis-MTS modifies non-desensitizing V666C receptors in other ways than current amplitude reduction. We measured the rate of receptor deactivation before and after the trap in M10M in the presence of CTZ and found no difference (Figure S5C-D, τ_pre-trap_ = 1.7 ms ± 0.2, τ_post-trap_ = 1.6 ± 0.2 ms, *n* = 8, *P* = 0.05 (paired randomisation test)). We considered the possibility that non-desensitizing receptors were silently modified by M10M. We tested this scenario with the following experiment: a patch with V666C receptors was first trapped in M10M and CTZ; CTZ was then washed-out the patch and freely desensitizing V666C receptors were exposed to M10M only (Figure S5A). If non-desensitizing V666C receptors had been silently modified by M10M, then a fraction of the receptors should have been protected resulting in the reduced sensitivity to further trapping by M10M. V666C receptors initially exposed to M10M in the presence of CTZ were modified to the same extent as naïve receptors by M10M once CTZ was unbound (I_peak_ post-trap / I_peak_ pre-trap for V666C initially trapped in CTZ: 0.23 ± 0.04, *n* = 6 and for V666C never trapped in CTZ: 0.30 ± 0.03, *n* = 16, *P* = 0.07; Figure S5B). Taken together, these results strongly suggest that non-desensitizing V666C receptors were not silently modified by M10M. Instead, the reduced inhibition of active receptors reflected state-dependent protection from modification.

### Auxiliary subunits do not alter the geometry of desensitized receptors

Trapping with bis-MTS cross-linkers so far indicated more conformational flexibility of the LBD layer in desensitized than activated AMPA receptors. However, synaptic AMPA receptors are rarely expressed alone, and are instead associated with various auxiliary proteins that re-define their kinetic properties (Schwenk et al., 2012; Jackson and Nicoll, 2011). We therefore wondered if the presence of auxiliary subunits, such as Stargazin (Stg), could affect conformational flexibility of AMPA receptors.

Recently, cryo-EM structures of AMPA receptors in complex with auxiliary subunits have been acquired in resting, active and desensitized states (Figure 1B) (Chen et al., 2017; Twomey et al., 2017a). These structures all predict reduced separation of V666C residues when compared to receptors without auxiliary proteins. For example, sulfhydryl groups of V666C residues on subunits A and C are 19 Å apart in the apparent active state of isolated receptors and 15 Å in activated receptors complexed with Stg (Figure 1C). In the desensitized state, the equivalent residues are 21 Å apart in homologous kainate receptors (PDB: 5KUF (Meyerson et al., 2016)) and 17 Å in desensitized GluA2 receptors associated with GSG1L auxiliary proteins (Figure 1C, PDB: 5VHZ (Twomey et al., 2017b)). If, indeed, auxiliary subunits keep the LBD layer more compact, we reasoned that their presence should also limit the effects of longer bis-MTS cross-linkers.

To test this hypothesis, we repeated the trapping experiments on complexes of AMPA receptors with Stg (Figure 6). GluA2 and Stg were co-expressed and association of complexes was assessed by measuring the ratio of kainate current over glutamate current (KA/Glu). The relative efficacy of the partial agonist kainate is known to be higher for GluA2-Stg complexes than for GluA2 alone, making it a good marker of GluA2-Stg association (Tomita et al., 2005; Shi et al., 2009). After establishing formation of the GluA2 V666C-Stg complexes in the patch, we proceeded with the trapping protocol that exposed complexes to glutamate and a bis-MTS cross-linker (1 μM) for 1 minute (Figure 6A-B), as described previously. No blocker of desensitization was added and the receptors were allowed to desensitize freely. Therefore, the crosslinking represents trapping across a mixture of active and desensitized states.

**Figure 6.**
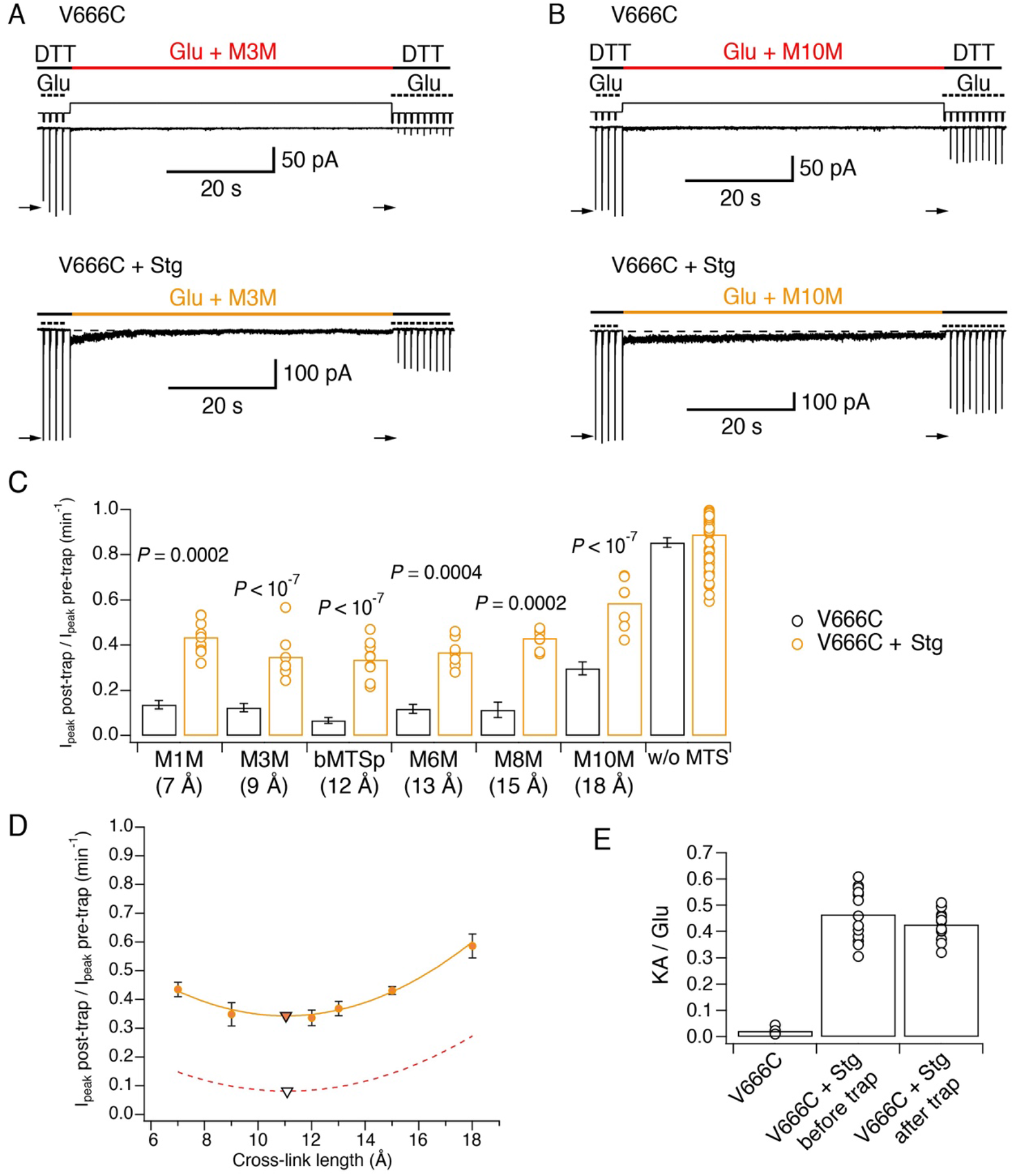
Stargazin attenuates effects of bis-MTS cross-linkers. (A) Current traces of V666C trapping in M3M, without (top) and with Stargazin (Stg; bottom). Legend is the same as in Figure 2, with a trapping pulse shown here in orange. (B) Same as in (A), but for M10M. (C) Trapping effects for V666C without (black) and with Stg (orange). Post- and pre-trap peak current was determined from the control pulses (arrows in (A) and (B)). Data without Stg are the same as in Figure 2D. *P* values compare the effects of the respective cross-linker with and without Stg. For statistics vs. w/o MTS and between cross-linkers, see Table S3. (D) Trapping profile of desensitizing V666C+Stg complexes. The data were fit with a parabola (orange line); the fit reaches minimum (orange triangle) at (11, 0.3). Trapping profile of desensitized receptors without Stg is shown as red, dashed line for comparison (minimum at (11, 0.1); black tringle). (E) The kainate/glutamate (KA/Glu) peak current ratio was determined for each patch before and after trapping with a bis-MTS (similar to the experimental design in Figure 3A-C with 1mM KA and 10 mM Glu in 1 mM DTT). V666C+Stg KA/Glu measurements before and after trap shown here are paired recordings, pooled for various bis-MTS cross-linkers.

The trapping results summarized in Figure 6C show that the presence of auxiliary subunit Stg apparently protected V666C receptors from cross-linking by bis-MTS. Indeed, following trapping, a robust response was preserved, and could not be overcome by longer trapping intervals (Figure S6A-B).

To test whether bis-MTS cross-linkers perhaps act on non-complexed V666C receptors only, without affecting V666C-Stg complexes, we measured the KA/Glu current ratio before and after the bis-MTS trap for a series of patches. We reasoned that if only non-complexed V666C receptors were being modified, the glutamate-activated current should reduce, but the kainate current (which is almost entirely carried by GluA2-Stg complexes) should not. Therefore, preferential trapping of non-complexed V666C mutants should lead to an increase in KA/Glu ratio. As shown in Figure 6E, the KA/Glu ratio was not affected (before trap: 0.46 ± 0.03; after trap: 0.43 ± 0.02, n = 13; *P* = 0.2, paired randomisation test), indicating V666C-Stg complexes were being modified by bis-MTS cross-linkers.

### Stargazin maintains active receptors in a compact arrangement

The trapping profile of V666C-Stg complexes (orange in Figure 6D) reflects the partial protection from trapping in the presence of Stg for all cross-linker lengths, but its overall shape is practically superposable onto the trapping profile of desensitized V666C receptors without Stg (red, dashed line in Figure 6D). Strikingly, the two curves reach their minimum at the same point of 11 Å (triangles in Figure 6D) and have identical curvature. This indistinguishable length dependence indicated that the trapping of V666C-Stg complexes came primarily from trapping desensitized receptors, and that the active complexes of V666C-Stg might be untouched by bis-MTS cross-linkers.

To test this hypothesis, we repeated the trapping protocol in the presence of CTZ, to block desensitization of V666C-Stg complexes. Two cross-linkers were tested in this condition: M1M, the shortest one, and M8M, which had the strongest trapping effect on active V666C receptors without Stg (Figure 5B-C). As predicted, neither bis-MTS cross-linker had any effect on activated GluA2-Stg complexes (M1M: *P* = 0.1, *n* = 9; M8M: *P* = 0.2, *n* = 9; paired recordings with and without bis-MTS) (Figure 7). Over very long exposures, M8M could induce irreversible inhibition, presumably from residual desensitization (Figure S6). This result confirmed that partial trapping observed for desensitizing V666C-Stg mainly derived from bis-MTS cross-linkers accessing desensitized complexes.

**Figure 7.**
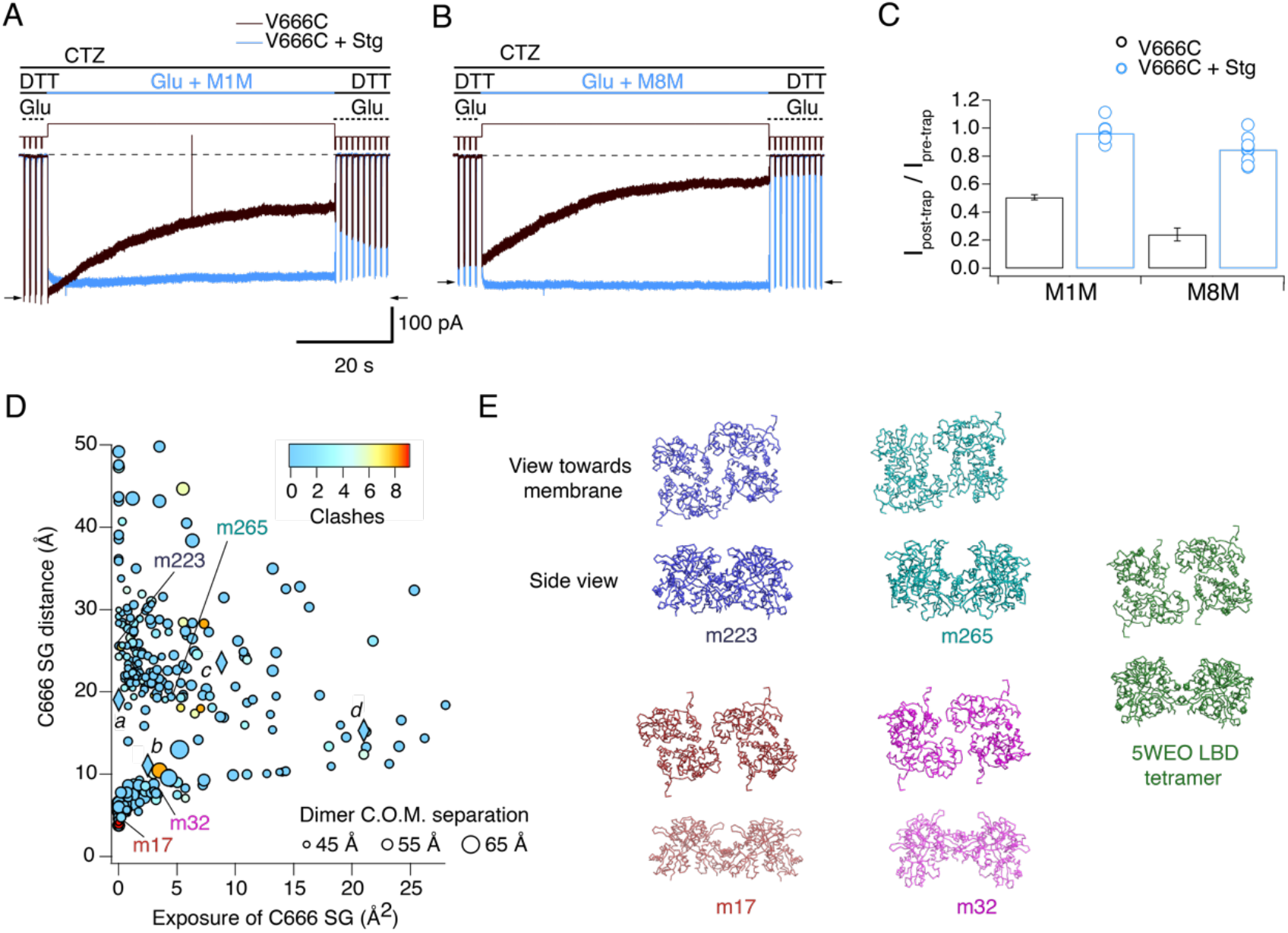
Stargazin blocks access to bis-MTS cross-linkers in the active state. (A) Traces showing trapping by M1M in the active state in the absence (black) and presence (blue) of Stargazin (Stg). Legend is the same as in Figure 2, with a trapping pulse shown here in blue. Desensitization was blocked by CTZ (100 μM) present throughout the experiment. Glutamate (Glu) was 10 mM and DTT 1mM. (B) Same as in (A), but for M8M cross-linker. (C) Summary of the trapping effects for M1M and M8M in the active state in the absence (black) and presence (blue) of Stg. Pre- and post-trap current was determined from control pulses as indicated by arrows. (D) Putative compact structures that could protect from bis MTS modification. Results of 268 runs of rigid body docking of LBD dimers against each other, to minimize Cys666 access in subunit A and mimic the protection from crosslinking. Results from known structures (5JEI, 4YU0 loose, 4YU0 tight and 5WEO, diamonds *a-d*) and 4 of the generated models – m17, m32, m223, m265 - segregate into two classes, similar to the loose (SG-SG distance between subunits < 10 Å) and tight arrangements (SG-SG distance > 20Å, C666 SG buried close to helix K) (Baranovic et al., 2016). The symbol size corresponds to the dimer centre of mass separation and the colour to the number of atom clashes (distance < 2.2 Å.) For reference, dimer centre of mass separations of known structures are: 5JEI, 44 Å; 4YU0 loose, 47 Å; 4YU0 tight, 47Å and 5WEO, 52 Å. Note that m17 is selected from a cluster of models with zero clashes (cyan), but adjacent in the graph to two other models with clashes (red circles). (E) LBD arrangements in four models marked on the graph in panel D and the original seed for each optimization, the glutamate bound LBD tetramer from 5WEO full-length GluA2 structure with Stg

We next wondered if the protection from trapping observed for active GluA2-Stg complexes could, at least partly, be explained by reduced accessibility of cysteine residues during exposure to bis-MTS, perhaps because the Cys666 residues are buried against other subunits or oriented such that the sulfhydryl groups are inaccessible for cross-linking. We mined possible conformations of the active LBD tetramer (PDB: 5WEO) using a coarse docking approach, allowing each active dimer to move in the membrane plane, and allowing rotation of one dimer with respect to the other around axes parallel and perpendicular to the active dimer interface (Figure 7D and E). Interestingly, docked structures with low Cys666 accessibility segregated into two classes, a loose arrangement where Cys666 were physically close but sterically hindering each other (with the centre of mass separation of the two active dimers comparable to 5WEO, e.g. models m17 and m32 in Figure 7D), and a tighter arrangement of the LBDs with V666C close to helix K (with the centre of mass separation of the two active dimers smaller than in 5WEO, e.g. m223 and m265 in Figure 7D), not unlike the structure of the V666C mutant LBD bound with fluorowillardine (5JEI) (Salazar et al., 2017). In that structure, additional shielding of the V666C side chain is afforded by its being shielded by neighbouring structural elements.

Thus, active GluA2-Stg complexes can avoid cross-linking not just by adopting more compact LBD arrangements, but also by preferring conformations which shield V666C SG groups.

## Discussion

Technical advances in cryo-electron microscopy have revolutionized the study of membrane proteins, and the supply of structural information is greater than ever before. However, as the catalogue of images swells, the need to relate their geometry to dynamics becomes ever more pressing. In case of AMPA receptors, full-length structures in complexes with various ligands and auxiliary subunits have been published. Here we have used a classical crosslinking approach allied to rapid perfusion to measure distances up to 18 Å in AMPA receptor extracellular domains across different functional states and in the time domain.

Bis-MTS cross-linkers have been used previously to aid identification of movements underlying state transitions in AMPA and NMDA receptors (Armstrong et al., 2006) (Tajima et al., 2016). Here, we report effects with a strong dependence on the target site, length of the cross-linker, functional state of the receptor and the presence or absence of auxiliary subunits. Since most bis-MTS cross-linkers are alkyl chains and hence flexible, we tested the bMTSp linker that includes a rigid benzene ring, but otherwise has a very similar length to M6M. The only difference we observed between these two cross-linkers was with K493C mutant, with M6M blocking desensitization more effectively than bMTSp, perhaps indicating an angle between the K493C side chains that was more easily accommodated by the flexible chain.

All bis-MTS cross-linkers inhibited the current, irrespective of the functional state. Whereas cross-linking of the desensitized states should result in current inhibition, cross-linking of an open channel would be expected to result in more active receptors. However, all inter-dimer constraints of the active state so far seem to be inhibitory (Plested and Mayer, 2009) (Lau et al., 2013; Baranovic et al., 2016; Yelshanskaya et al., 2016). Whereas the zinc and disulphide bridges, previously used to cross-link active LBD dimers, only form under strict geometrical requirements, the same cannot be said for bis-MTS cross-linkers. The cross-linkers are flexible and therefore do not restrict the trapped LBD tetramer to a single geometry. The inhibitory effect of the crosslinkers is not universal – at the active intra-dimer interfaces we could potentiate receptors as shown previously (Armstrong et al., 2006). It appears that crosslinks between dimer pairs, or indeed any restriction between the active dimers, leads to a decrease in activity. The mechanism behind this inhibition remains frustratingly unclear.

In Figure 8A-B, the trapping profiles of V666C receptors with the available structural models of GluA2 in the equivalent condition are compared. The shallow trapping profile of desensitized AMPA receptors indicates a structural ensemble of conformations broadly in agreement with multiple desensitized states (Meyerson et al., 2014) (Robert and Howe, 2003). Even though the LBD dimer-dimer separation observed in the model (>18 Å, EMDB: 2688) was not tested with the bis-MTS cross-linker of the corresponding length, extrapolation of the trapping profile indicates such conformations are available to desensitized AMPA receptors. However, the longest cross-linker (18 Å) was the slowest one to act on desensitized receptors, indicating that this conformation, and more ‘dilated’ ones, are not readily available for cross-linking, but slowly get populated during prolonged exposures to agonist. The longest cross-linker, M10M, was also the slowest one to act in the active state. There, the most readily accessible separation of V666C residues, in terms of strength and speed of trapping, was at 15 Å (green in Figure 8B), with 19 Å separation, predicted by the structural model (PDB: 4UQ6), taking longer to populate. Notably, ‘dilations’ of extracellular domains were also seen in antagonist-bound NMDA receptors (Zhu et al., 2016).

**Figure 8.**
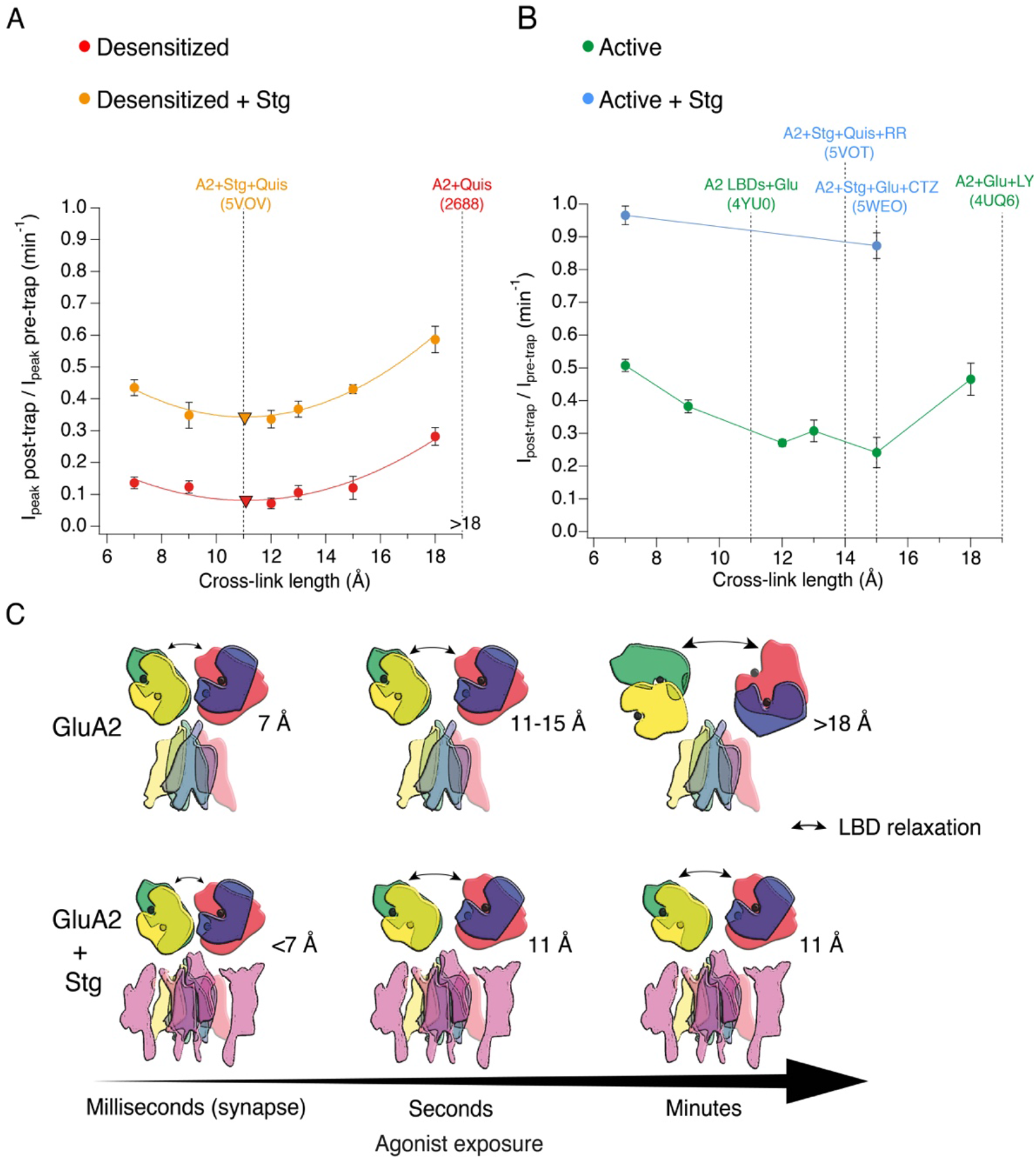
The presence of Stargazin and short agonist exposures keep the agonist-bound LBD layer compact. (A) Trapping profiles for V666C in desensitized states, without (red) and with (orange) Stargazin (Stg). Solid lines are fit parabola, with local minima indicated as triangles. For desensitized condition without Stg, minimum is reached at (11, 0.1) and with Stg at (11, 0.3). The right handside of the x-axis is marked as >18 Å, as the resolution of the only available desensitized structure without auxiliary subunits is too low to measure residue distances. Vertical, dashed, black lines indicate V666C sulfhydryl separation in the respective structure (PDB accession codes in brackets). Names of structures are colour-coded same as the trapping profiles. (B) Same as in (A), but for active state without (green) and with (orange) Stg. 4YU0 is a structure of soluble, isolated LBDs. Quis stands for quisqualate (full agonist), RR for (R, R)-2b (Kaae et al., 2007) and LY for LY451646 (both desensitization blockers). (C) Schematic summarizing the bis-MTS cross-linking results. Colour-code same as in Figure 1; ATDs are omitted. Black spheres represent bound glutamate. Upper row: in the absence of Stg, GluA2 receptors access ‘dilated’ LBD conformation (black arrows) upon long exposures to agonist. Bottom row: presence of Stg limits ‘dilation’ of the LBD layer even at longer agonist exposures.

The presence of Stg lead to universal attenuation of the cross-linking effect, for all bis-MTS lengths and both functional states, desensitized and active (orange and blue in Figure 8A and B, respectively). The longest crosslinker (M10M) produced the least trapping, having reduced trapping potency even following long exposures (~3 min). Thus, the LBD layer could not enter a widely-splayed, dilated form for majority of these receptors. This result seems to chime with structural and FRET experiments (Shaikh et al., 2016), where the presence of Stg made the LBD and ATD layer more compact. In the desensitized state, the shape of the trapping profile is the same, with or without Stg (Figure 8A), with both parabola reaching a minimum point at 11 Å (triangles in Figure 8A). This is in excellent agreement with the structure of desensitized GluA2-Stg complex (PDB: 5VOV). In the active state, our results deviate from the available full-length active structures. Both short and long (7 or 15 Å) bis-MTS reagents failed to inhibit currents in the first minute of trapping, indicating more compact LBD arrangements over this timescale than obtained in the structures.

The protection from trapping in our cross-linking experiments most likely has multiple origins. Although “protection” could result from a persistent long-distance separation, outside the range of the crosslinkers, several observations and common sense speak against this possibility. First, we previously showed that active receptors (glutamate + CTZ) could be trapped by zinc bridges in compact arrangements (Baranovic et al., 2016). Second, the long distance must be maintained throughout the exposure, because any transit between compact and dilated arrangements must pass through intermediate separations, allowing crosslinkers to span the gap. Third, even the most dilated structures are in the range of crosslinker lengths that we used. Fourth, the mixed trapping condition for GluA2-Stg complexes in the absence of CTZ apparently supports desensitized state trapping over a wide range of geometries but no additional active state trapping. We cannot discount specific, state-dependent protection from crosslinking, produced by a unique, closed-cleft LBD conformation provoked by Stg in the active state, but it seems to us unlikely. We reasoned that protection from crosslinking could, at least partly, be accounted for by reduced accessibility of cysteine residues during exposure to bis-MTS. Docking data in Figure 7D-E show that conformations which reduce the accessibility of the V666C side chains are indeed accessible to the active LBDs in the presence of Stg, with some of them acquiring more compact arrangements than in the current structural models of the full-length receptor in the active state (Chen et al., 2017; Twomey et al., 2017a).

No matter how avid and complete trapping is, a possible source of receptor activity after trapping is the activity of free (non-crosslinked) subunits (Figure S7). In all our experiments, subunits *A* and *C* are trapped whereas *B* and *D* are not. In GluA2 receptors without Stg, even with desensitization blocked, the activity of only two subunits (e.g. *B* and *D*) produces low conductance openings with short open times (Rosenmund et al., 1998). But the presence of Stg (Coombs et al., 2017) (Zhang et al., 2014) increases the current carried by receptors with one or two active subunits. In our experiments, Stg had an indefatigable effect of maintaining receptor activity in the limit of long exposures to bis-MTS. The effect of Stg was inordinately large, with a maximum effect of the crosslinking leaving approximately half the receptor activity unscathed. Can the two non-crosslinked subunits (*B* and *D*) produce this level of activity? A simple back-of-the-envelope calculation suggests this effect is larger than expected. Single channel recordings of GluA2 with Stg reveal that the conductance from two subunits, with desensitization blocked, should be on average about 40% but with *P*_open_ < 1 (Coombs et al., 2017). Therefore, residual activity of the non-crosslinked subunits is higher than expected, unless the B and D subunits have a predominant role in driving channel gating (as proposed from structural studies (Sobolevsky et al., 2009).

We presume that the basal compactness of the LBD layer is related to observation that longer cross-linkers have slower trapping rates, indicating slow adoption of ‘dilated’ conformations. Higher bis-MTS concentrations sped up the reactions accordingly, while leaving the relative order of trapping rates intact. Alkylthiosulfonates are generally distinguished by their extremely rapid reactivity in mild conditions, selectivity for cysteinyl sulfhydryl groups, general reversibility upon addition of thiols such as DTT and their ability to effect quantitative and complete conversion to the disulphide without applying a large excess of reagent (Kenyon and Bruice, 1977). Our reaction rates are close to the maximum expected (10^5^ M^−1^s^−1^; (Liu et al., 1997)).

Physiological activation and desensitization of AMPA receptors takes place on a millisecond timescale, and we monitored this process wherever possible during our experiments. Generally, the remaining current responses were not altered following bis-MTS exposures. The fast gating contrasts to desensitized states trapped by bis-MTS cross-linkers that take tens of minutes to recover. Though we necessarily worked at low bis-MTS concentrations to avoid confounding effects like chaining and non-specific modification, mandating slow trapping, the slow recovery from trapping is striking. It seems reasonable to assume that the degree of stabilization by different crosslinkers is similar, and that the difference in stability comes from the states themselves. This idea is supported by the cut-off that we observe – for crosslinkers longer than 10 Å, crosslinking in desensitized states is irreversible over 10 minutes (also in 5-fold higher concentration of the reducing agent). It is possible that upon cross-linking, V666C residues adopt conformations that make them inaccessible to the reducing agent, but this would then have to be highly specific for bis-MTS reagents longer than 9 Å and state-specific (due to much faster recovery rates of active than desensitized receptors after trapping in M1M or M3M). In addition, in reducing conditions, AMPA receptors recover from disulphide crosslinking at the same sites in hundreds of milliseconds (Lau et al., 2013; Salazar et al., 2017). Assuming that the length of the bis-MTS cross-linkers reflects level of structural rearrangements and following the trend plotted in Figure 4H, this suggests that AMPA receptors exposed to brief glutamate transients at synapses are unlikely to undergo extreme conformational changes, due to their complexation with auxiliary subunits. Considering that all of the experiments presented here were done with over-expressed receptors and auxiliary subunit Stg, it is possible that V666C-Stg complexes had variable stoichiometry of association, depending on the expression level. This could impact how Stg affects conformational dynamics of the receptors, but a detailed investigation of this is beyond the scope of this work. We note, however, that the presence of two copies of auxiliary subunit GSG1L were sufficient to keep the LBD layer compact in structural experiments (Twomey et al., 2017b).

Long agonist exposures of minutes to hours are standard in structural biology experiments and could, thus, contribute to the prevalence of more ‘relaxed’ conformations in full-length structures without auxiliary subunits (Figure 8C). However, long exposures to agonists are at odds with synaptic conditions where AMPA receptors see glutamate on a millisecond timescale, before it is actively cleared by transporters (Clements, 1996). Furthermore, any large structural rearrangements of the extracellular domains would need to be accommodated by the crowded synaptic environment (High et al., 2015; Tao et al., 2018) and potential presynaptic interaction partners (Saglietti et al., 2007) (Elegheert et al., 2016). When in complex with auxiliary subunits, no functional state of AMPA receptors necessitates large domain movements, and compact arrangements of the extracellular layer can sustain the gating process. We conclude that extracellular domains of synaptic AMPA receptors are unlikely to undergo large structural rearrangements during synaptic transmission and instead work in a fairly compact conformational regime, unless faced with long exposures to glutamate in pathological conditions.

## Acknowledgements

We thank Clarissa Eibl and Mark Mayer for comments on the manuscript. This work was funded by the ERC Grant “Gluactive” (647895) and the NeuroCure Cluster of Excellence (DFG EXC-257). A.J.R.P. is a Heisenberg Professor of the DFG (Project number 323514590).

## Author Contributions

J.B. and A.J.R.P. designed experiments, J.B. performed experiments, A.J.R.P. wrote PYTHON code and did the docking computations. J.B. and A.J.R.P. analysed data and wrote the manuscript.

## Experimental Procedures

### Molecular Biology

In all experiments, the unedited (Q586) GluA2flip version of rat GluA2 gene was expressed from the pRK5 vector. Amino acid numbering refers to the mature receptor assuming a signal peptide of 21 amino acids in length. As a marker of transfection, eGFP was expressed from the same vector, downstream from an internal ribosomal entry sequence (IRES). Mouse Stargazin gene (a kind gift from Susumu Tomita) was expressed from a separate pRK8 vector containing IRES-dsRed (Carbone and Plested, 2016). All mutations were introduced by overlap PCR and confirmed by double-stranded sequencing.

### Cell Culture and Transfection

GluA2 constructs were expressed transiently in HEK293 cells using calcium-phosphate precipitation or PEI method as described previously (Baranovic et al., 2016; Riva et al., 2017). HEK293 cells were obtained from the Leibniz Forschungsinstitut DSMZ (Deutsche Sammlung von Mikroorganismen und Zellkulturen GmbH, Germany) ACC no. 305 (RRID: CVCL_0045) and tested negative for mycoplasma. Cells were maintained in MEM Eagle medium (PAN-Biotech GmbH, Aidenbach, Germany) supplemented with 10% (v/v) fetal bovine serum and antibiotics (penicillin (100 U/mL) and streptomycin (0.1 mg/mL; PAN-Biotech).

For transfections, 2-3 μg of DNA was transfected per 35 mm dish and cells were washed after 6-8 hours. Recordings were performed 24-72 hours after the transfection at room temperature. For transfections with Stargazin, Stargazin DNA was co-transfected with GluA2 DNA at 2:1 mass ratio and after the transfection, cell medium was supplemented with 40 μM NBQX to reduce Stargazin-induced cytotoxicity.

### Solutions

Chemicals were obtained from Sigma Aldrich (Munich, Germany), Abcam plc (Cambridge, UK) and Hello Bio (Bristol, UK). MTS compounds were obtained from Toronto Research Chemicals (North York, Canada).

The internal (pipette) solution for recordings without Stargazin contained (mM): 115 NaCl, 1 MgCl_2_, 0.5 CaCl_2_, 10 NaF, 5 Na_4_BAPTA, 10 Na_2_ATP, 5 HEPES, titrated to pH 7.3 with NaOH. For recordings with Stargazin, the internal solution was slightly modified: 120 NaCl, 0.5 CaCl_2_, 10 NaF, 5 Na_4_BAPTA, 5 HEPES and 0.05 spermine, pH 7.3. The external recording solution in all experiments contained (mM): 150 NaCl, 0.1 MgCl_2_, 0.1 CaCl_2_ and 5 HEPES, pH 7.3. Different drugs were added to the external solution as needed. Glutamate was always applied at 10 mM and DL-dithiothreitol (DTT) at 1 mM. Cyclothiazide (CTZ) and kainate (KA) were thawed from stock solutions on the day of the experiment. Final CTZ and KA concentrations in all experiments were 100 μM and 1 mM, respectively.

All MTS compounds were obtained as powder. Whereas monofunctional MTS reagents are known to be highly reactive and unstable in aqueous solutions (Kenyon and Bruice, 1977), this information is lacking for bifunctional cross-linkers used in this study. Hence, we took special care to minimize exposure of bis-MTS compounds to oxidizing (aqueous) solutions (Takatsuka and Nikaido, 2010). The powder was dissolved in DMSO, aliquoted and kept on ice on the day of the experiment. Once a stable patch recording was obtained, an aliquot was dissolved in external solution to a final concentration of 1 μM and applied to the patch. This way, aqueous bis-MTS solutions were on average 2-3 minutes old at the moment of application. Each MTS stock was tested with GluA2 wild-type receptors and K493C receptors as negative and positive controls, respectively. The final MTS concentration of 1 μM was chosen based on previous work (Sobolevsky et al., 2003; Yelshansky et al., 2004). We avoided higher concentrations of MTS compounds due to potential cross-reactivity and chaining effects.

### Patch clamp electrophysiology

Ligands and drugs were applied to outside-out patches via a custom made 4-barrel glass (VitroCom, USA) mounted to a linear piezo–electric wafer (PiezoMove P-601.4, PI, Germany) (Lau et al., 2013). Two barrels were perfused with control solutions and the third barrel with the trapping solution, as described below. All patches were voltage clamped at −40 mV unless stated otherwise. Currents were low-pass filtered at 10 kHz (−3 dB cut-off, eight-pole Bessel filter) using an Axopatch200B amplifier (Molecular Devices, U.S.A.) and acquired with AxographX software (Axograph Scientific, Australia, RRID:SCR_014284) at 20 kHz sampling rate via Instrutech ITC-18 digitizer (HEKA, Germany). Current traces were digitally filtered at 1 kHz (low-pass) for presentation in figures.

To assess the effect of different bifunctional cross-linkers on AMPA receptors, the receptors were exposed to the cross-linker (1 *μ*M) for 1 minute. Before and after this trapping exposure, the current in the patch was tested with control pulses that contained only 10 mM glutamate, without the cross-linker and in the presence of DTT (1 mM) as a reducing agent (Figure 2A-C). Four control pulses before application of the cross-linker provided a measure of the patch current before any exposure to the cross-linker. Accordingly, (up to thirty) control pulses recorded after the MTS application, were used to assess any changes in the patch current imparted by the cross-linker treatment.

For recordings of GluA2 receptors co-expressed with auxiliary subunit Stargazin, care must be taken that GluA2 receptors are indeed associating with Stargazin. One strategy to minimize the presence of lone V666C receptors relies on the relief of spermine (polyamine) block at positive voltages imparted by complexation with Stargazin (Carbone and Plested, 2016). Although we have included spermine in the pipette solution and measured relieve of block for each patch, we did not perform recordings at positive voltages, as the currents were not stable enough during minutes-long trapping protocols. Instead, a change in kainate efficacy was used as a marker of GluA2-Stargazin association as described in the text.

### Analysis

Trapping effects were quantified as the ratio of the average current after the trap (determined from the 2^nd^ post-trap control pulse) and average current before the trap (determined from the 4 pre-trap control pulses; arrows in Figure 2A-C):

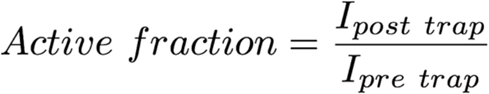

In case of desensitizing receptors, peak current was measured and in the case of non-desensitizing receptors, steady-state current.

The trapping time of each cross-linker was determined from cumulative exposures to a bis-MTS of up to 6 minutes (6 repetitions of the trapping protocol). After each application, current reduction was determined with respect to the initial current in the patch, before any trap. The resulting current decay was described by a monoexponential fit in Igor Pro (v7.06, Wavemetrics, Lake Oswego, Oregon, U.S.A., RRID:SCR_000325).

To determine the rate of recovery from trapping by MTS cross-linkers, the number of post-trap control pulses was increased until full recovery was attained. An envelope of post-trap peak current responses was then created in Igor Pro and fit with a monoexponential. This approach was possible only for faster recovery rates, on the time scale of seconds, such as recovery of desensitized V666C receptors from trapping with M1M (Figure 2B and 4D) and recovery of active V666C receptors from trapping with M1M and M3M (Figure 5A). With longer bis-MTS cross-linkers, the recovery time increased from seconds to minutes, making direct measurements of the recovery time from post-trap control pulses impractical. Instead, the experimental design was re-adjusted to allow measurements of long recovery times as described in Figure 4A-C. In brief, peak current in the patch was initially recorded with 100 ms jumps into glutamate in control conditions until it stabilized. Then, a trapping protocol was performed as described above, with control pulses before and after the trap. After the trapping protocol, the current in the patch was again monitored, for about 10 minutes, with fast control jumps into glutamate in order to follow any potential recovery of the peak current. In this time period, desensitized V666C receptors managed to recover only from trapping by M1M and M3M (Figure 4D-G), in which case the recovery was fit with a monoexponential.

Trapping profiles (Figure 2G and 6D) were fit with a parabola in Igor Pro:

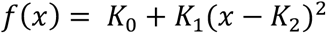

where *K*_1_ defines the curvature, *K*_0_ the minimum effect and *K*_2_ the x value at the minimum. Data points were weighted by the standard error of the mean for the fit.

### Computational docking

To investigate structures of the LBD tetramer that could preclude trapping by blocking access to the Cys666 SG moiety, we treated each dimer as a rigid body and subjected them to rotations and translations in the membrane plane. Python scripts (available at github.com/aplested/cystance) were written as a glue for molecular manipulations in PyMOL molecular manipulations and CCP4 (RRID:SCR_007255) functions to measure geometry and exposure of the Cysteine (AREAIMOL, NCONT (Winn et al., 2011)). For each run, trial arrangements that reduced Cys666 SG accessibility in subunit *A* whilst also keeping the dimers in close proximity (with minimal atom clashes) and maintaining physiologically plausible in-plane linker arrangements were retained as seeds for subsequent rounds, and the step size was reduced. Trial arrangements with more than 10 atom clashes (< 2.2 Å) were rejected. No refinement was done to eliminate spurious clashes from flexible surface residues. The optimization was ended when no further improvement was possible. Each search lasted about 5-10 minutes on a 2017 Macbook Pro.

All *P* values were determined by non–parametric randomisation test (non–paired, unless stated otherwise), using at least 10^5^ iterations (DC-Stats suite: https://github.com/aplested/DC-Stats). Bars in graphs indicate mean and error bars SEM.

Structural models and related measurements were visualized and measured in PyMOL (v2.0, RRID:SCR_000305) (The PyMOL Molecular Graphics System, Version 2.0 Schrödinger, LLC). The length of bis-MTS cross-linkers was measured in ChemDraw Professional (PerkinElmer, U.S.A.).

